# The orphan nuclear receptor NR4A3 is dispensable for resident memory CD8^+^ T cell generation

**DOI:** 10.1101/2024.09.09.612076

**Authors:** Livia Odagiu, Salix Boulet, Dave Maurice De Sousa, Jean-François Daudelin, Nathalie Labrecque

**Affiliations:** Montreal Clinical Research Institute, Montreal, Qc, Canada, H2W 1R7; Maisonneuve-Rosemont Hospital Research Center, Montreal, Qc, Canada, H1T 2M4; Département de microbiologie, infectiologie et immunologie, Université de Montréal, Qc, Canada, H3C 3J7; Département de médecine, Université de Montréal, Qc, Canada, H3C 3J7

## Abstract

Different memory CD8^+^ T cell subsets are generated following acute responses: central, effector and resident (Trm). CD8^+^ Trm cells established residency at the sites of infection and provide an efficient and rapid frontline defense against re-infection. The NR4A family members (NR4A1, NR4A2 and NR4A3) of orphan nuclear receptor are transiently expressed following TCR signaling and NR4As were shown to influence CD8+ T cell response. Interestingly, *Nr4a1*, *Nr4a2* and *Nr4a3* have been reported to be transcribed by CD8^+^ Trm cells. In absence of NR4A1, less CD8^+^ Trm cells are present in the liver, lungs, small intestine intra-epithelial lymphocytes (IELs) and Peyer’s patches. NR4A2 was shown to play a role in the generation of small intestine IEL CD8^+^ Trm cells. However, evidence is still lacking for the contribution of NR4A3 during CD8^+^ Tm cell differentiation. In this study, we evaluated the role of NR4A3 in the differentiation and maintenance of CD8^+^ Trm cells. Our data demonstrate that in contrast to the other family members NR4A1 and NR4A2, NR4A3 is dispensable for the generation of CD8^+^ Trm cells in both epithelial and non-epithelial sites.

## Introduction

Different memory CD8^+^ T (Tm) cell subsets are generated following acute responses: central (Tcm), effector (Tem) and resident (Trm). Tcm cells circulate within lymphoid organs and have the capacity to re-expand following antigen re-encounter while Tem cells have immediate effector functions and circulate within the blood to patrol the organism^1,2^. Trm cells develop from KLRG1^lo^ precursors that seed the infected tissues where they establish residency^3^. They provide an efficient and rapid frontline defense against re-infection by contributing directly to the elimination of the infectious agent and by recruiting circulating Tem and Tcm cells to the infected sites^4–7^.

CD8^+^ Trm cells can be further divided into the ones that seed epithelial tissues, such as skin, and non-epithelial tissues, such as liver. CD8^+^ Trm cells resident at epithelial sites are characterized by expression of CD69 and CD103 while the non-epithelial Trm cells do not express CD103^8^. Depending on the tissues in which CD8^+^ Trm cells reside their dependency on TGF-β will differ^8,9^. Skin Trm cells are dependent on TGF-β for their generation while liver Trm cells are not^8^. The presence of TGF-β signals in epithelial sites regulates CD103 expression, explaining why Trm cells at non-epithelial sites are CD103 negative^8^.

The expression of NR4A family members (NR4A1, NR4A2 and NR4A3) of orphan nuclear receptors is rapidly induced following TCR signaling and are considered as being good reporter of T cells receiving a TCR signal^10–12^. They have been reported to control several aspects of T cell development and response^13^. During acute response, we have shown that NR4A3 deficiency increases the generation of MPECs and central memory CD8^+^ T cells and improves effector functions^14^.

*Nr4a1*, *Nr4a2* and *Nr4a3* have been reported to be transcribed by CD8^+^ Trm cells^3,15–18^. In absence of NR4A1, less CD8^+^ Trm cells are present in the liver, lungs, small intestine intra-epithelial lymphocytes (IELs) and Peyer’s patches^18^. Using a genetic screen, Milner et al identified NR4A1, NR4A2 and NR4A3 as possible modulators of small intestine IEL CD8^+^ Trm differentiation in although their role was not further validated^15^. NR4A2 was shown to play a role in the generation of CD8^+^ Trm cells as knockdown of its expression in CD8^+^ T cells decreased the generation of small intestine IEL CD8^+^ Trm cells^16^. However, evidence is still lacking for the contribution of NR4A3 during CD8^+^ Tm cell differentiation. In this study, we evaluated the role of NR4A3 in the differentiation and maintenance of CD8^+^ Trm cells. Our data demonstrate that in contrast to the other family members NR4A1 and NR4A2, NR4A3 is dispensable for the generation of CD8^+^ Trm cells in both epithelial and non-epithelial sites.

## Results

### The generation of resident memory CD8^+^ T cells in secondary lymphoid organs is not affected by deficiency of NR4A3

To evaluate the role of NR4A3 in the differentiation and maintenance of CD8^+^ Trm cells in different tissues, we adoptively transferred *Nr4a3*^+/+^ or *Nr4a3*^-/-^ OT-I T cells (CD45.2^+^) in competition with wild-type OT-I T cells (CD45.1^+^) from B6.SJL mice into CD45.1^+^CD45.2^+^ congenic mice followed by infection one day later with a recombinant strain of LCMV encoding the ovalbumin antigen (LCMV-OVA) (Fig.1A). The response of CD8^+^ T cells was followed over time using CD45.1 and CD45.2 staining as illustrated in Fig.1B, where *Nr4a3*^+/+^ or *Nr4a3*^-/-^ OT-I T cells are identified as CD45.2^+^ while the competitor wild-type OT-I from B6.SJL are CD45.1^+^. Although we injected the OT-I cells and their competitors at a 1:1 ratio, we always observed a stronger response at day 8 post-infection (Fig. 1B). A similar observation is made when *Nr4a3*^+/+^ OT-I are in competition with OT-I from B6.SJL at memory time point in the spleen (Fig. 2A). As reported by us^14^, *Nr4a3*^-/-^ OT-I T cells generates more memory CD8^+^ T cells than their wild-type counterpart in the spleen at day 30 post-infection (Fig. 2A) with an increase in Tcm cells as shown using CD62L staining (Fig. 2B). The enhanced generation of Tcm was accompanied by a decrease in KLRG1^+^ memory CD8^+^ T cells in absence of NR4A3 (Fig. 2B). Similar observations were made in the mesenteric lymph nodes (LNs) (Fig. 2C-D).

**Figure 1.**
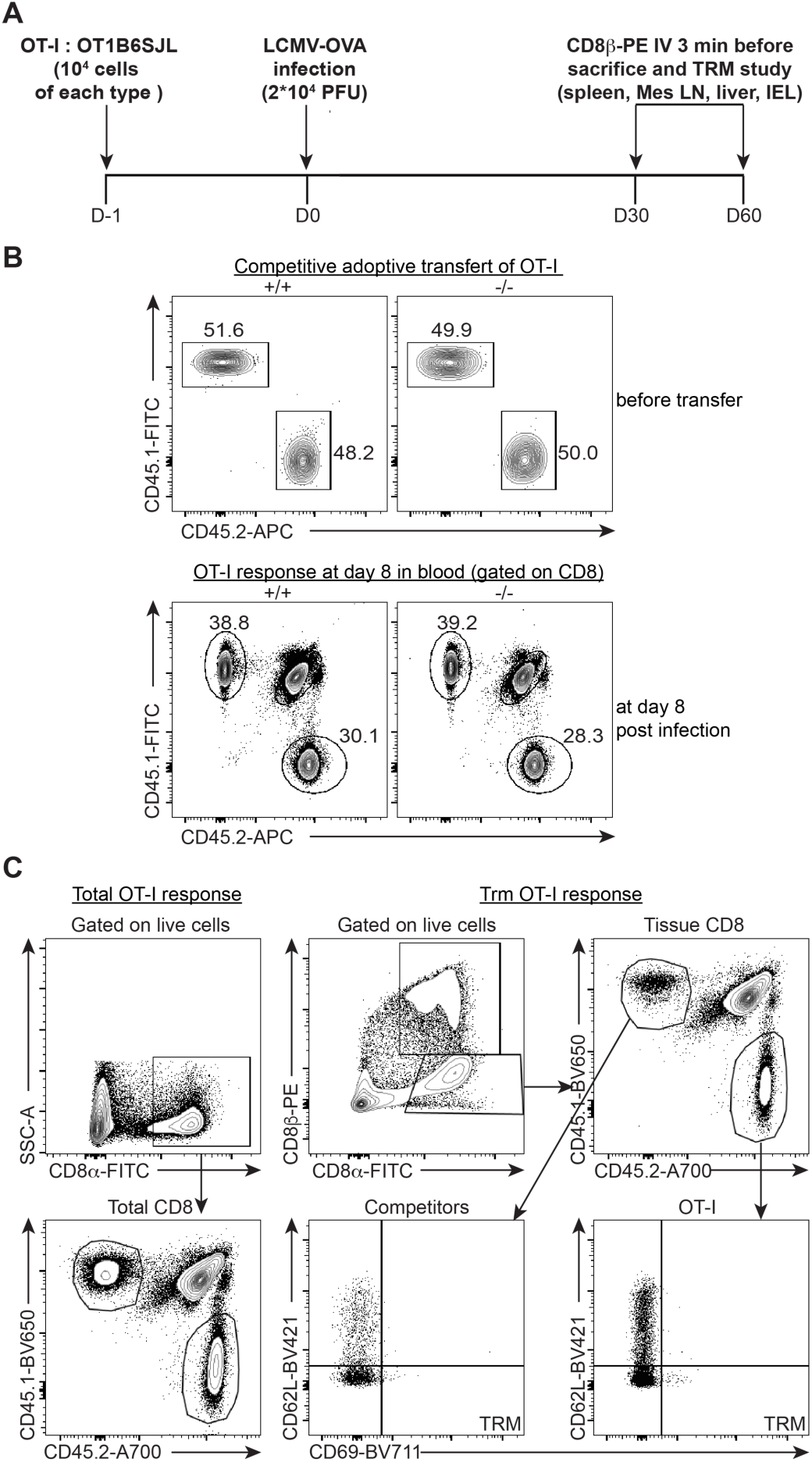
Experimental design for the study of the role of NR4A3 during CD8^+^ Trm cell differentiation. **A.** *Nr4a3*^+/+^ and *Nr4a3*^-/-^ OT-I (CD45.2^+^) T cells were adoptively transferred in competition with wild-type OT-I B6.SJL competitors (CD45.1^+^) into CD45.1^+^/CD45.2^+^ hosts followed by infection with LCMV-OVA one day later. **B**. *Nr4a3*^+/+^ or *Nr4a3*^-/-^ OT-I (CD45.2^+^) and OT-I B6.SJL cells ratio prior the adoptive transfer into CD45.1^+^/CD45.2^+^ recipient mice (top) and their response in the blood at day 8 post infection with LCMV-OVA (bottom). **C.** Gating strategy to evaluate the total OT-I memory CD8^+^ T cell response and to identify *Nr4a3^+/+^* and *Nr4a3^-/-^* OT-I cells (CD45.2^+^) versus to their OT-I B6.SJL competitors (CD45.1^+^) CD8^+^ Trm cells.

**Figure 2.**
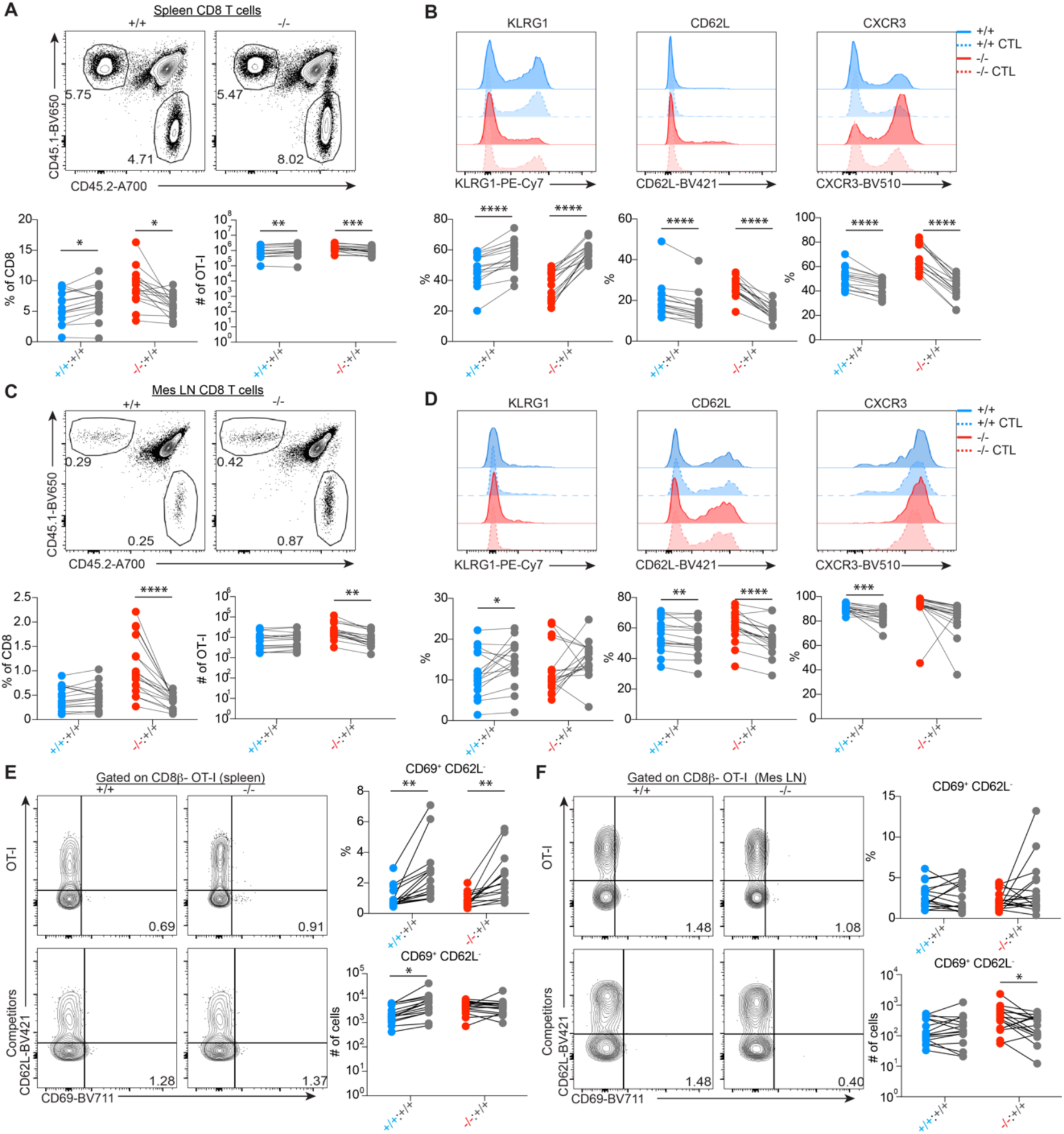
NR4A3 deficiency does not affect CD8^+^ Trm cell differentiation in secondary lymphoid organs. Response of OT-I (CD45.2^+^, *Nr4a3*^+/+^ or *Nr4a3*^-/-^) and their competitors OT-I B6.SJL (CD45.1^+^) at day 30 post-infection with LCMV-OVA in the spleen (**A**) and mesenteric lymph nodes (Mes LN) (**C**). KLRG1, CD62L and CXCR3 expression by OT-I T cells at day 30 post-infection with LCMV-OVA in the spleen (**B**) and Mes LN (**D**). (**E-F**) Quantification of CD8^+^ Trm cell generation in the spleen (**E**) and Mes LN (**F**) at day 30 post-infection with LCMV-OVA. CD8^+^ Trm cells were analysed following i.v. injection of anti-CD8β. CD8^+^ Trm cells were identified as CD8β^-^CD69^+^CD62L^-^. Each competitor pair was compared by a paired Student t-test. \**P*<0.05, \*\**P*<0.01, \*\*\**P*<0.001, \*\*\*\**P*<0.0001.

Resident memory CD8^+^ T cells are generated in the spleen and LNs following LCMV infection^17,19,20^. We therefore evaluated whether NR4A3 controls the generation of CD8^+^ Trm cells in these organs. As shown in Fig. 2E-F, we did not observe significant differences in the generation of CD8^+^ Trm cells (CD69^+^CD62L^-^) between *Nr4a3*^+/+^ and *Nr4a3*^-/-^ OT-I T cells in the spleen and mesenteric LNs at day 30 post-infection.

Our results demonstrate that NR4A3 is dispensable for the generation of CD8^+^ Trm cells in secondary lymphoid organs. We then tested whether NR4A3 is required for the maintenance of CD8^+^ Trm cells, a likely possibility as they transcribed *Nr4a1/2/3* genes^3,15,16^. As shown in Figure 3A-D, at day 60 post-infection with LCMV-OVA, we observed more OT-CD8^+^ memory T cells in lymphoid organs with an increase proportion of CD62L^+^ Tcm cells when OT-I T cells lack NR4A3 expression. However, we did not observed differences in Trm cells (Fig. 3E-F) further indicating that NR4A3 is not important for their maintenance in secondary lymphoid organs.

**Figure 3.**
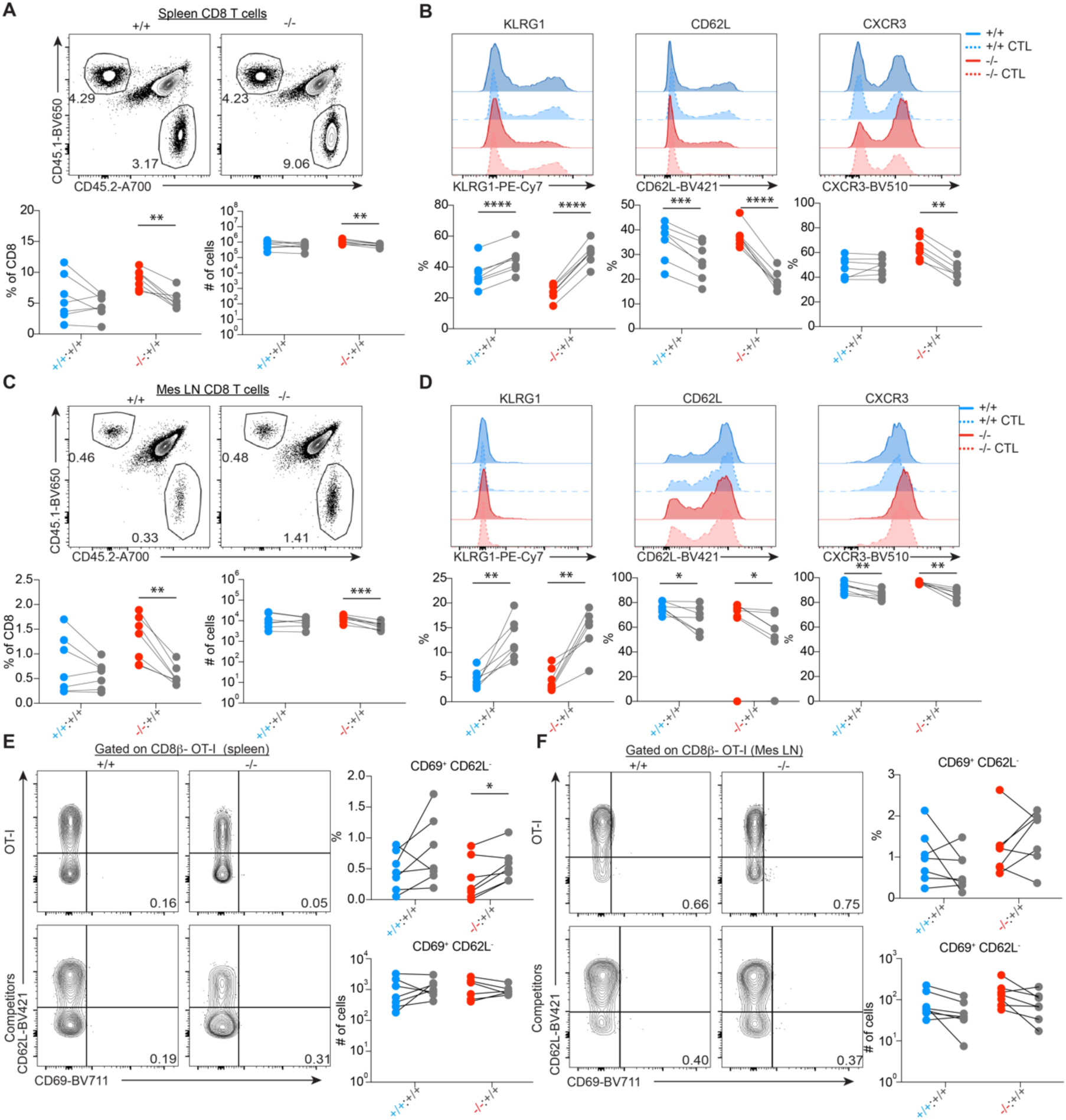
NR4A3 expression is not required for the maintenance of CD8^+^ Trm cells in secondary lymphoid organs. Response of OT-I (CD45.2^+^, *Nr4a3*^+/+^ or *Nr4a3*^-/-^) and their competitors OT-I B6.SJL (CD45.1^+^) at day 60 post-infection with LCMV-OVA in the spleen (**A**) and mesenteric lymph nodes (Mes LN) (**C**). KLRG1, CD62L and CXCR3 expression by OT-I T cells at day 60 post-infection with LCMV-OVA in the spleen (**B**) and Mes LN (**D**). (**E-F**) Quantification of CD8^+^ Trm cells in the spleen (**E**) and Mes LN (**F**) at day 60 post-infection with LCMV-OVA. CD8^+^ Trm cells were analyzed following i.v. injection of anti-CD8β. CD8^+^ Trm cells were identified as CD8β^-^CD69^+^CD62L^-^. Each competitor pair was compared by a paired Student t-test. \**P*<0.05, \*\**P*<0.01, \*\*\**P*<0.001, \*\*\*\**P*<0.0001.

### The generation of resident memory CD8^+^ T cells is not influenced by NR4A3 at non epithelial sites

It is well recognized that molecular events controlling the generation and maintenance of CD8^+^ Trm cells in different tissues are different. The signals controlling the generation and maintenance of CD8^+^ Trm cells are distinct whether they formed at non epithelial sites (non-barrier tissue) such as the liver when compared to epithelial sites (skin, intestine). Therefore, we evaluated whether NR4A3 was controlling the generation and maintenance of CD8^+^ Trm cells in the liver, a non-epithelial site. To evaluate this, we took advantage of the fact that LCMV infects a broad range of tissues and thus generates CD8^+^ Trm cells at epithelial and non-epithelial sites. In contrast to what we have observed in the secondary lymphoid organs, we did not observe an increase in the percentage and number of OT-I CD8^+^ memory T cells in the liver when comparing *Nr4a3*^+/+^ and *Nr4a3*^-/-^ T cells (Fig. 4A-B). This might reflect the fact that NR4A3 deficiency acts on Tcm cells, which are not colonizing the liver. Similar results were obtained at day 60 (Fig.4C-D). We then evaluated whether NR4A3 affects the differentiation of memory CD8^+^ T cells into Trm cells. At day 30 post-infection, we did not observe any difference between *Nr4a3*^+/+^ and *Nr4a3*^-/-^ OT-I T cells in their ability to generate CD8^+^ Trm cells as shown with no difference in the proportion and number of CD69^+^CD62L^-^ OT-I T cells (Fig. 4E). The maintenance of CD8^+^ Trm cells was not affected by NR4A3 deficiency (Fig. 4F) and we noted a tendency for better maintenance of the *Nr4a3*^-/-^ OT-I Trm cells in the liver. Therefore, NR4A3 expression is dispensable for the generation and maintenance of CD8^+^ Trm cells in the liver.

**Figure 4.**
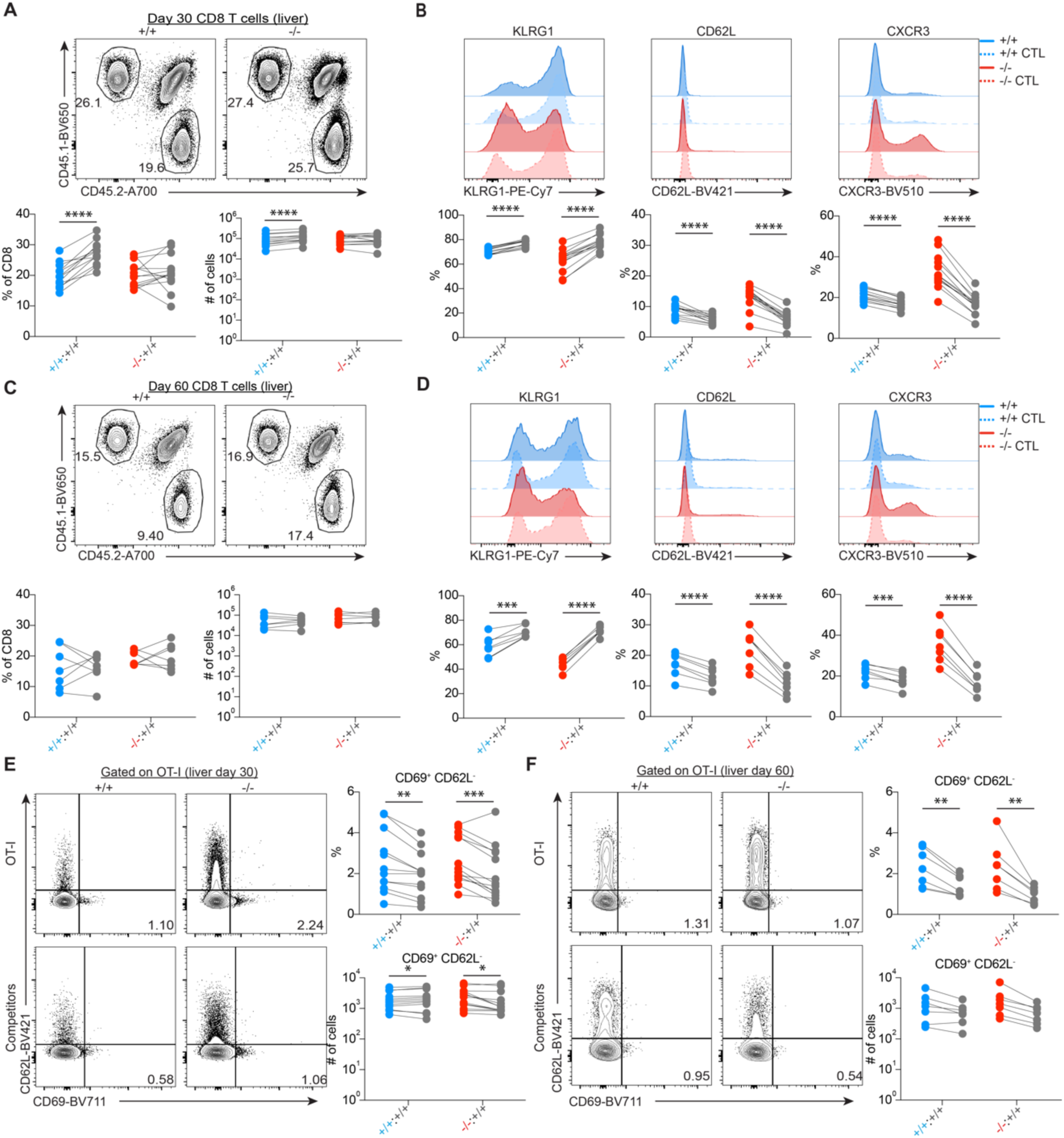
NR4A3 deficiency does not impact liver CD8^+^ Trm cell differentiation. Response of OT-I (CD45.2^+^, *Nr4a3*^+/+^ or *Nr4a3*^-/-^) and their competitors OT-I B6.SJL (CD45.1^+^) at day 30 (**A**) and day 60 (**C**) post-infection with LCMV-OVA in the liver. KLRG1, CD62L and CXCR3 expression by OT-I T cells at day 30 (**B**) and 60 (**D**) post-infection with LCMV-OVA in the liver. (**E-F**) Quantification of CD8^+^ Trm cell generation in the liver at day 30 (**E**) and 60 (**F**) post-infection with LCMV-OVA. CD8^+^ Trm cells were identified as CD69^+^CD62L^-^. Each competitor pair was compared by a paired Student t-test. \**P*<0.05, \*\**P*<0.01, \*\*\**P*<0.001, \*\*\*\**P*<0.0001.

**Figure 5.**
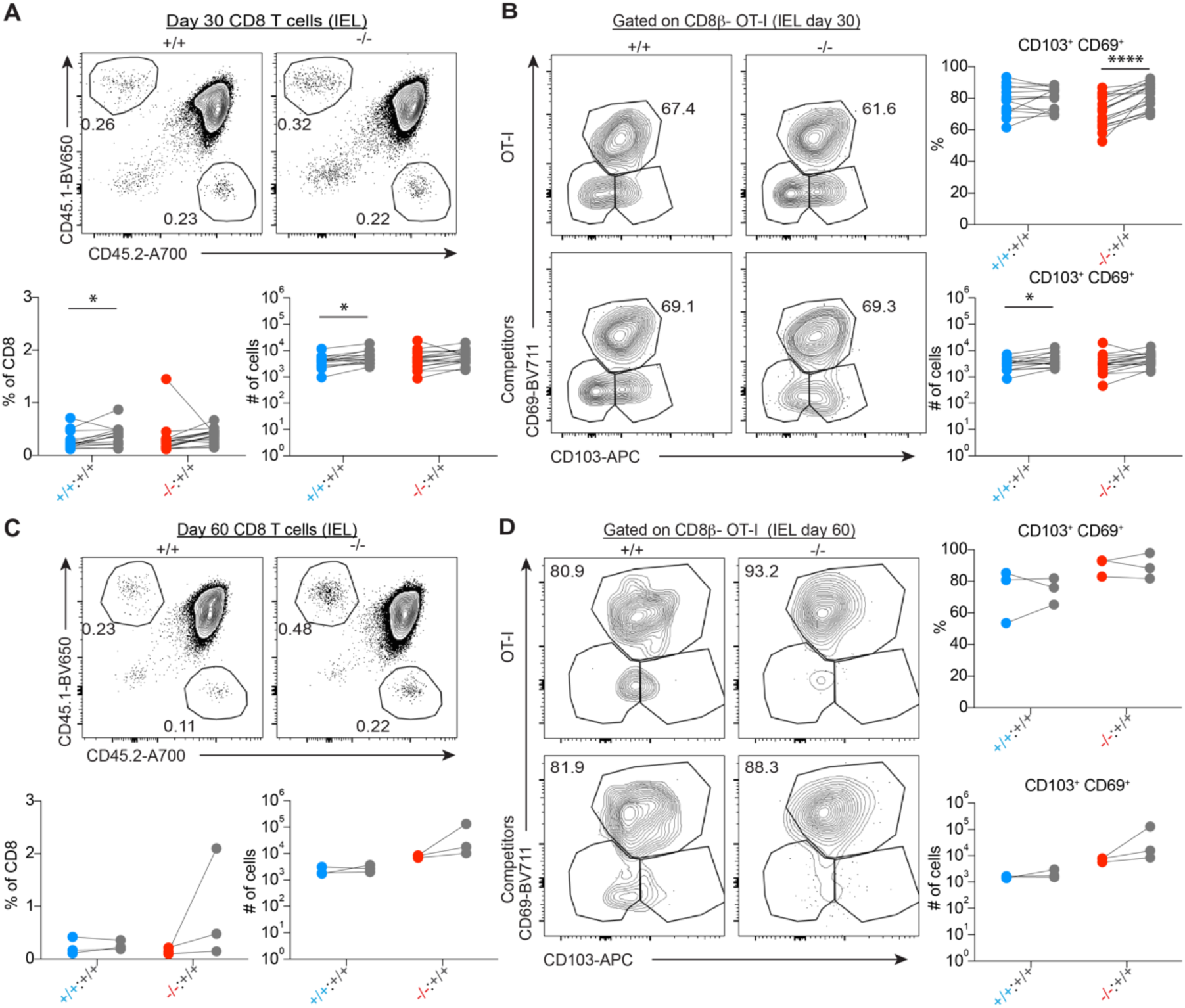
NR4A3 is dispensable for CD8^+^ Trm cell differentiation in the small intestine. Response of OT-I (CD45.2^+^, *Nr4a3*^+/+^ or *Nr4a3*^-/-^) and their competitors OT-I B6.SJL (CD45.1^+^) at day 30 (**A**) and 60 (**C**) post-infection with LCMV-OVA in the small intestine IELs. Quantification of CD8^+^ Trm IELs at day 30 (**B**) and 60 (**D**) post-LCMV-OVA infection. CD8^+^ Trm cells were analysed after i.v. injection of anti-CD8β. CD8^+^ Trm IELs were identified as CD8β^-^CD69^+^CD103^+^. Each competitor pair was compared by a paired Student t-test. \**P*<0.05, \*\**P*<0.01, \*\*\**P*<0.001, \*\*\*\**P*<0.0001.

### NR4A3 is dispensable for the generation of resident memory CD8^+^ T cells in epithelial tissues

The requirement for the generation of epithelial CD8^+^ Trm cells being different than for non-barrier sites, we investigated whether NR4A3 influences the generation if intra-epithelial CD8^+^ Trm cells in the intestine. To identify CD8^+^ Trm cells within IELs, we injected anti-CD8β-PE antibody i.v. 3 minutes before sacrifice allowing to distinguish circulating (CD8β-PE^+^) from tissue-resident (CD8β-PE^-^) T cells (Fig. 1C). As for the liver, we did not observe increase OT-I memory CD8^+^ T cell generation in the IELs and no difference in cells with a Trm cell phenotype (CD69^+^CD103^+^ and CD69^+^CD103^-^) at day 30 and 60 post-infection with LCMV-OVA. This demonstrates that NR4A3 is dispensable for the generation of CD8^+^ Trm cells in both epithelial and non-epithelial sites.

## Discussion

The transcription of *Nr4a1*, *Nr4a2* and *Nr4a3* at the steady state by CD8^+^ Trm cell ^3,15,16^ suggests that they contribute to the generation or maintenance of CD8^+^ Trm cells. This has been firmly demonstrated for NR4A1 and NR4A2^15,16,18^. In this manuscript, we evaluated the role of NR4A3 in the generation and maintenance of CD8^+^ Trm cells in different tissues using LCMV infection. Our results demonstrate that NR4A3 is dispensable for the differentiation and maintenance of CD8^+^ Trm cells in secondary lymphoid organs, liver and small intestine.

The dispensable role for NR4A3 contrasts with the one that was reported for NR4A1 and NR4A2^15,16,18^. This may indicate that the other family members have redundant role with NR4A3. This is surprising as we have shown that NR4A3 deficiency alone is sufficient to improve the generation of CD8^+^ Tcm cells^14^. Therefore, NR4A3 has unique function during CD8^+^ T cell response when compared to NR4A1 and NR4A2. Interestingly, CD8^+^ Tcm cells do not transcribe the *Nr4a1/2/3* genes, which indicates that NR4A3 acts before memory CD8^+^ T cells are generated, in agreement from our previous report of an early programming role of NR4A3 during acute CD8^+^T cell response^14^. Furthermore, the enhancement of Tcm cell differentiation in absence of NR4A3 does not lead to a reduction of CD8^+^ Trm cell generation, indicating that NR4A3 does not control a Tcm versus Trm cell fate choice. Furthermore, the impact of NR4A3 deficiency on the Trm cell differentiation may be masked by the fact that *Nr4a3*^-/-^ CD8^+^ T cells generate more KLRG1 negative cells, the precursor of CD8^+^ Trm cells^3^, when compared to their wild-type counterpart. Such an impact on the generation of KLRG1 negative Trm precursors was not observed with *Nr4a1*^-/-^ CD8^+^ T cells^21^ (unpublished observations).

Another possible explanation for a lack of effect of NR4A3 for the generation of CD8^+^ Trm cells may relate to its level and kinetic of induction. Indeed, it was reported that *Nr4a3* transcription requires stronger TCR signals than for *Nr4a1*^11^. Therefore, at the steady state it is possible that less *Nr4a3* is transcribed than the other members which then might be insufficient to contribute to CD8^+^ Trm cell differentiation. Furthermore, in all the studies reporting expression of the *Nr4a* genes, only transcriptional readouts were used, it is thus possible that the level of NR4A3 protein might be insufficient to affect CD8^+^ Trm cell generation.

The differential effect of NR4A family members on distinct stages of the CD8^+^ T cell response suggests that although the three members recognize the same DNA motif they regulate different set of genes. Further studies are required to identify the target genes of each family member. One possible mechanism by which different members act differently could be the consequence that only NR4A1 and NR4A2 can heterodimerize with RXR, another nuclear receptor^22,23^.

In conclusion, our study demonstrates that in contrast to NR4A1 and NR4A2, NR4A3 is dispensable for the generation of CD8^+^ Trm cell generation while it dampens the generation of CD8^+^ Tcm cells.

## Material and methods

### Mice

OT-I (*Rag1^-/-^* CD45.2) ^24^, *Nr4a3^-/-^* ^25^ and *Nr4a3^+/+^*, B6.SJL and C57BL/6 mice were bred at the Maisonneuve-Rosemont Hospital Research Center animal facility. CD45.1.2 mice were obtained from F1 cross of B6.SJL and C57BL/6 mice. OT-I (*Rag1^-/-^* CD45.2) mice were crossed with B6.SJL (CD45.1) to obtain OT-I/B6.SJL mice (*Rag1^-/-^* CD45.1). *Nr4a3^-/-^* mouse stain was a kind gift from Dr. Orla M. Conneely^25^. OT-I *Nr4a3^+/+^* (*Rag1^-/-^* CD45.2) and OT-I *Nr4a3^-/-^*(*Rag1^-/-^* CD45.2) mice were obtained by crossing *Nr4a3^-/-^*mice (backcrossed to C57BL/6 for at least 10 generations) with OT-I mice. All the mice were housed in an animal facility pathogen-free environment and treated in accordance with the CCAC (Canadian Council on Animal Care) guidelines.

### Adoptive transfer and LCMV-OVA infection

OT-I cells were isolated from lymph nodes of OT-I *Nr4a3^+/+^,* OT-I *Nr4a3^-/-^* and OT-I/B6.SJL mice. 10^4^ OT-I CD8^+^ T cells (CD45.2^+^) were mixed in a ratio of 1:1 with OT-I/B6.SJL CD8^+^ T cells (CD45.1^+^) and adoptively transferred i.v. into CD45.1.2 (CD45.1^+^ CD45.2^+^) recipient mice. The following day adoptively transferred mice were infected with 2 x 10^4^ PFU of LCMV Armstrong encoding OVA (LCMV-OVA) by i.v. injection. LCMV-OVA virus strain was a kind gift from Juan C. de la Torre (The Scripps Research Institute, La Jolla, CA, USA)^26^. LCMV-OVA virus was produced in the cell culture supernatant of infected L929 fibroblast cell line (cultured in MEM supplemented with 5 % heat inactivated Nu serum) and the virus titer was determined in MC57G fibroblasts^27^.

### Isolation of CD8^+^ Trm cells from different tissues

An i.v. injection with 3µg of CD8β-PE (clone YTS 156.7.7 Biolegend, cat #126607) was done 3 min before the mouse sacrifice and the organs (spleens, liver, mesenteric LN and small intestines) were collected in complete RPMI media^14^ and kept on ice until processing. Spleen and LNs were dissociated between frosted glass slides and spleen red blood cells were lysed by a 5 min RT incubation with 0.83% hypotonic NH_4_Cl (Biobasic, CAS #12125-02-9) solution. The liver was chopped into small pieces (2 to 3 mm) and crushed with a syringe plunger before the collagenase D (1mg/ml in complete RPMI; Sigma life science, #11088882001) digestion 10 min at 37°C. The liver cell suspension was then passed through a 100µm cell strainer (Fisher, # 0877119), followed by a red blood cells lysis. Liver lymphocytes were then isolated at the interphase of a 40 on 80% Percoll (Cytiva 17-0891-01) gradient centrifugation (2000 rpm, 20 min at RT without brake). For IEL isolation, Peyer’s patches, the fat and the conjunctive tissue were all removed from small intestine and then they were opened to rinse out the content. Cleaned intestine were cut into 2 to 3 cm pieces and shaked at 300 rpm 20 min at 37°C in a 5mM EDTA (Corning, 46-034-Cl) and 0,145mg/ml DTT complete RPMI media (3% Nu serum). Intestine pieces were then vigorously vortexed for 30 sec and passed through a strainer and this process was repeated two times by collecting intestine pieces in 2mM EDTA containing RPMI (0% Nu serum). The IEL cell suspension was then filtered through a 70µm cell strainer (Fisher, 087712) and the collected cells were then resuspended in complete RPMI (3% Nu serum) and filtered through a 40µm cell strainer (Fisher, 352340). IEL cells were then resuspended in 30% Percoll and centrifuged at 1700 rpm 20 min at RT without brake. After Percoll centrifugation the IEL cell pellet was washed with complete RPMI.

### Flow cytometry

Freshly extracted cell suspension from different organs were washed with PBS 1X prior to a 10 min incubation with Fc Block (clone 2.4G2, Leinco Technologies, C381-1.0mg) and the viability dye (Zombie NIR, Biolegend, 423106) at RT. The extracellular staining was then performed 20 min on ice as previously described^14,28,29^. Flow cytometry data acquisition was done on BD LSRFortessa X-20 and BD LSR II from BD Biosciences. Cytometry data were analyzed with FlowJo software (Tree Star). The antibodies used in this study are CD8α-FITC (clone 53-6.7, Biolegend, cat #100706), CD45.1-BV650 (clone A20, Biolegend, cat #110736), CD45.2-A700 (clone 104, Biolegend, cat #109822), CD45.1-FITC (clone A20, Biolegend, cat # 110706), CD45.2-APC (clone 104, Biolegend, cat #109814), CD69-Biotin (clone H1.2F3, Biolegend, cat #104504), Streptavidin-BV711 (Biolegend, cat#405241), CD62L-BV421 (clone MEL-14, Biolegend, cat #104436), KLRG1-PE-Cy7 (clone 2F1/KLRG1, Biolegend, cat #138416), CXCR3-BV510 (clone CXCR3-173, Biolegend, cat #126528), CD103-APC (clone 2E7, Biolegend, cat #121414).

### Statistical analyses

A paired Student t-test comparison was done for each competitor pair (OT-I *Nr4a3^+/+^* or OT-I *Nr4a3^-/-^* with OT-I/B6.SJL). The statistical analysis was performed using Prism software (GraphPad Software). Data are presented as individual samples. \**P* < 0.05, \*\**P* < 0.01, \*\*\**P* < 0.001, \*\*\*\**P* < 0.0001 were considered statistically significant.

## Acknowledgements

We thank Dr O. Conneely for the kind gift of *Nr4a3^-/-^* mice and Juan C. de la Torre for providing LCMV-OVA. **Funding:** The Canadian Institutes of Health Research supported this work by a grant (PJT-168910) to N. Labrecque. L. Odagiu was supported by a studentship from the Fonds de la recherche Québec-Santé. **Author contribution:** L.O., S.B. and N.L. designed the experiments. L.O., S.B., D.M.D.S., J.-F.D. performed the experiments. L.O. analysed the data and prepared the figures. N.L. and L.O. wrote the paper. **Competing interests**: The authors declare that they have no competing interests.

